# SR9009 Regulates Macrophage Polarization via the NR1D1/NF-κB Axis to Ameliorate Obesity Associated Ulcerative Colitis

**DOI:** 10.1101/2025.06.01.657286

**Authors:** Zhi Xue Zheng, Xuan Cai, Ya Qi Liu, Jing Tao Bi

## Abstract

**Objective:** The incidence of ulcerative colitis (UC) and obesity has risen in recent years, potentially linked through metabolic dysregulation and chronic inflammation. The nuclear receptor NR1D1 is pivotal in regulating circadian rhythms and plays a significant role in inflammation and metabolism. This study investigates the therapeutic effects and mechanisms of the NR1D1 agonist SR9009 on obesity-related UC.

**Methods:** We established a mouse model of obesity-induced UC utilizing a high-fat diet alongside dextran sulfate sodium (DSS). 32 male C57BL/6 mice were divided into four groups: control (DZ), high-fat diet (GZ), obesity associated UC model group (UC), and SR9009 intervention group (JD), with eight mice each. We evaluated body weight, blood lipids, colonic tissue alterations, IL-1β, IL-18, macrophage polarization, and NR1D1 expression levels.

**Results:** Mice in the UC group demonstrated significantly elevated body weight, spleen index, TG, CHOL and inflammatory markers (*P*<0.01). Pathological scores of colonic tissues increased markedly (*P*=0.000), with a rise in M1 macrophages (CD68⁺) and a decline in M2 macrophages (CD206⁺) (*P*<0.001). NR1D1 expression was notably downregulated (P<0.01). Post-SR9009 intervention, the JD group showed significantly reduced serum TG and CHOL levels (*P*=0.001, 0.011), IL-1β and IL-18 (P<0.001), improved colonic pathology (*P*=0.000), a decrease in M1 macrophages, an increase in M2 macrophages, and an enhanced M1/M2 ratio (P<0.001).

**Conclusion:** SR9009 mitigates intestinal inflammation in obesity associated UC by activating NR1D1, inhibiting the NF-κB pathway, and modulating macrophage polarization (suppressing M1 and enhancing M2). These findings propose a novel strategy for targeting NR1D1 in the treatment of obesity-related ulcerative colitis.

## Introduction

Ulcerative Colitis (UC) is a chronic, recurrent non-specific inflammatory bowel disease primarily affecting the colon and rectum, impacting both the mucosal and submucosal layers. Its clinical manifestations are diverse, typically encompassing diarrhea, mucus-purulent stools, and abdominal pain, with some patients experiencing systemic symptoms [1–2]. The etiology of UC remains elusive, with prior studies suggesting associations with environmental factors, genetic predispositions, infections, gut microbiota, immune responses, and psychological factors [3–7]. In China, the prevalence of UC is increasing, becoming a significant public health concern, likely linked to changes in dietary habits and lifestyles [8–10]. Furthermore, numerous studies have established a correlation between obesity and UC, indicating that high-fat diet consumption and reduced physical activity may precipitate obesity, leading to a systemic low-grade inflammatory state that influences UC progression. Additionally, obesity may alter gut microbiota diversity, exacerbating inflammatory responses and worsening UC [11–14].

The biological clock system within organisms serves as a vital regulatory mechanism, primarily responsible for controlling behavioral and physiological circadian rhythms. The alternation of these rhythms broadly participates in regulating life activities, influencing physiological functions such as metabolic processes and behavioral manifestations while responding to environmental stimuli, including temperature, feeding, and chemical factors [15–17]. Numerous studies have demonstrated that disruptions in circadian rhythms are associated with inflammatory diseases, metabolic disorders, and psychiatric conditions [18–21]. The rhythmic oscillation of the biological clock relies on autoregulatory transcription/translation feedback loops involving multiple genes, including CLOCK, BMAL1, and NPAS2. The nuclear receptor subfamily 1 group D member 1 (NR1D1) is a critical molecule in regulating the expression of circadian rhythm genes and in linking the central and peripheral clocks [22–24]. As an additional feedback loop for circadian rhythms, activating NR1D1 enhances the binding of its protein to the BMAL1 promoter, directly inhibiting BMAL1 transcription, thus completing the auxiliary feedback process. Through this mechanism, NR1D1 integrates circadian rhythms with other biological functions, regulating metabolic gene expression and subsequently influencing glucose metabolism, lipid metabolism, and inflammatory responses [25,26]. Circadian rhythms are essential for maintaining the intestinal epithelial barrier; mutations in clock genes or changes in environmental signals may disrupt circadian rhythms, leading to increased secretion of inflammatory cytokines and activation of inflammation-related signaling pathways, ultimately exacerbating colitis. Significant changes in circadian rhythm-related gene expression have been observed in patients with UC, with reductions noted not only in intestinal tissues but also in circulating monocytes [27–29]. Due to the unclear pathogenesis of UC and the limited treatment options available, the exploration of effective therapies is of utmost importance [30,31]. This study aims to investigate the relationship between the circadian rhythm molecule NR1D1 and obesity-associated UC, providing insights for future treatment strategies for such diseases and others related to circadian rhythm disruptions.

## Materials and Methods

### 1.1 Animal Models

Thirty-two male C57BL/6 mice, weighing 20±2g and aged 7 weeks, were procured from SBF (Beijing) Biotechnology Co., Ltd. The mice were housed under a 12-hour light/dark cycle to simulate natural lighting conditions, maintained at 25°C with 60% humidity, and provided with ad libitum access to water and food. This study was approved by the Animal Ethics Committee of Beijing Jishuitan Hospital.

### 1.2 Key Instruments and Reagents

An automatic biochemical analyzer was acquired from Zhongshan Xinxing Medical Equipment Technology Co., Ltd. (Model: XR 220PLUS); a microplate reader from BIO-RAD (Model: 680); a vortex mixer from Haimen Qilin Bell Instrument Manufacturing Co., Ltd. (Model: GL-88B); a centrifuge from Hunan Hengnuo Instrument Equipment Co., Ltd. (Model: 2-16R); a UV spectrophotometer from Biochrom, UK (Model: 550); an electrophoresis apparatus from Beijing Baijing Biotechnology Co., Ltd. (Model: BG-subMIDI); an SDS-PAGE electrophoresis system from BIO-RAD (Model: mini-PROTEAN® Tetra Cell with miniTrans-Blot® Module And PowerPac™ Universal Power Supply); a flow cytometer from Changzhou Bideke Biotechnology Co., Ltd. (Model: BeamCyte-1026); and an electron microscope from Leica (Model: DM3000). Dextran sulfate sodium (DSS) was sourced from Sigma (Batch No. 48267); ELISA kits (Catalog No: MD120111), hematoxylin-eosin (HE) staining kits (Catalog No: MD911467, MD911477), and mouse IL-1 (Catalog No: MD163564), IL-18 (Catalog No: MD163564, MD120111) were obtained from Beijing Baiou Science Biomedical Technology Co., Ltd.; NR1D1 antibodies were acquired from Proteintech, USA (Catalog No: 14506-1-AP); NR1D1 agonist SR9009 was purchased from MCE (Catalog No: HY-16989).

### 1.3 Establishment of Animal Models

In this study, 32 male C57BL/6 mice were selected, weighing 20±2g and aged 7 weeks, acclimated for one week, and randomly divided into four groups of eight mice each:

1. Control group (DZ): Normal diet for 14 weeks, followed by 5 days of intraperitoneal saline injection (100μl) until sacrifice;
2. High-fat diet group (GZ): Fed a high-fat diet for 12 weeks, validated by weight gain, followed by 5 days of saline injection (100μl) until sacrifice at 14 weeks;
3. Obesity with ulcerative colitis model group (UC): Fed a high-fat diet for 12 weeks, then given a 2.5% DSS solution in drinking water for 8 days (high-fat + DSS for 8 days), with saline injections (100μl) administered from days 9 to 13, and sacrifice on day 14;
4. Agonist group (JD): Fed a high-fat diet for 12 weeks, then given a 2.5% DSS solution for 8 days, with SR9009 administered at a dose of 50 mg/kg (volume 100μl) from days 9 to 13, and sacrifice on day 14.

### 1.4 Sample Collection and Processing

Mice from each group were anesthetized with 1% pentobarbital sodium, at a volume of 0.0042ml/g. Blood was collected via ocular puncture (greater than 1ml) and divided into two portions: one for total blood stored at -80°C for subsequent PCR detection, and the other for serum after standing for one hour, then centrifuged at 3000 rpm for 15 minutes at 4°C. All mice underwent collection of spleen, liver, and thymus tissues, which were weighed, and organ indices calculated. Colonic tissues were sectioned into four parts, each approximately 1cm in size, designated for pathological detection (hematoxylin-eosin staining), PCR analysis, Western blotting (WB) analysis, and flow cytometry.

### 1.5 Pathological Analysis of Colonic Tissues

Colonic tissues were stained following HE kit instructions, and microscopic examination was performed to assess structural integrity. Pathological scoring focused on epithelial tissue destruction and inflammatory infiltration, with the sum forming the histological activity index (HAI) (Table 1) [32].

**Table 1.**
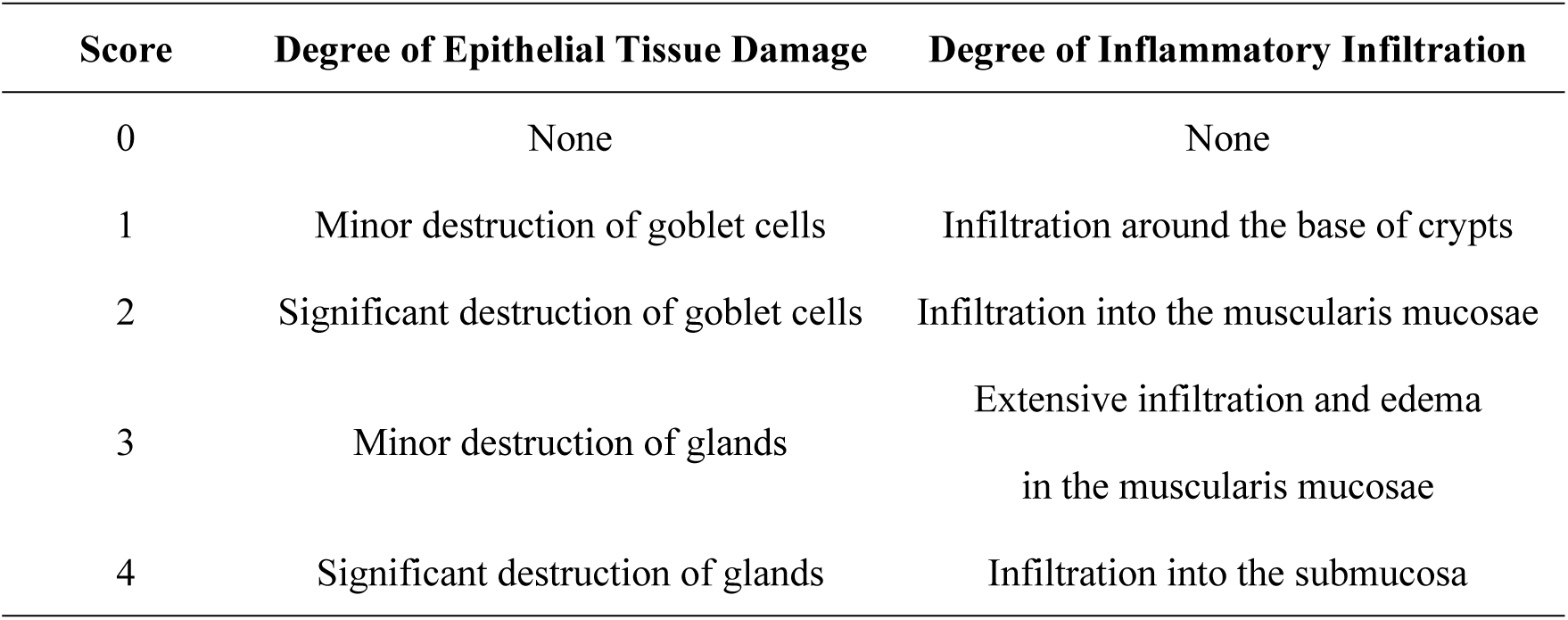
Pathological Scoring Criteria for HAI in Mouse Colonic Tissu.

### 1.6 Serum Biochemistry and Inflammatory Factor Detection

Serum was extracted and analyzed promptly according to kit instructions to determine TG and CHOL levels, and IL-1β and IL-18 levels were measured using ELISA kits.

### 1.7 Detection of NR1D1 and Inflammatory Factors in Serum and Colonic Tissues

Appropriate colonic tissue samples were homogenized using a tissue grinder. Total RNA was extracted according to kit guidelines. Using EXONGEN’s reverse transcription reagent ExonScript RT SuperMix with dsDNase, cDNA synthesis was performed. Real-time quantitative PCR was utilized to amplify target genes. Primers are detailed in Table 2, with the following reaction conditions: pre-denaturation at 95°C for 5 minutes; amplification in three steps at 95°C for 10 seconds, 58°C for 20 seconds, and 72°C for 20 seconds; and melt curve analysis at 95°C for 15 seconds, 65°C for 60 seconds, and 95°C for 15 seconds. GAPDH was used as a reference gene, with relative expression calculated using the 2-△△Ct method. Serum samples were collected, total RNA extracted per kit instructions, and IL-1, IL-18, and NR1D1 relative expression levels were assessed using real-time quantitative PCR.

**Table 2.**
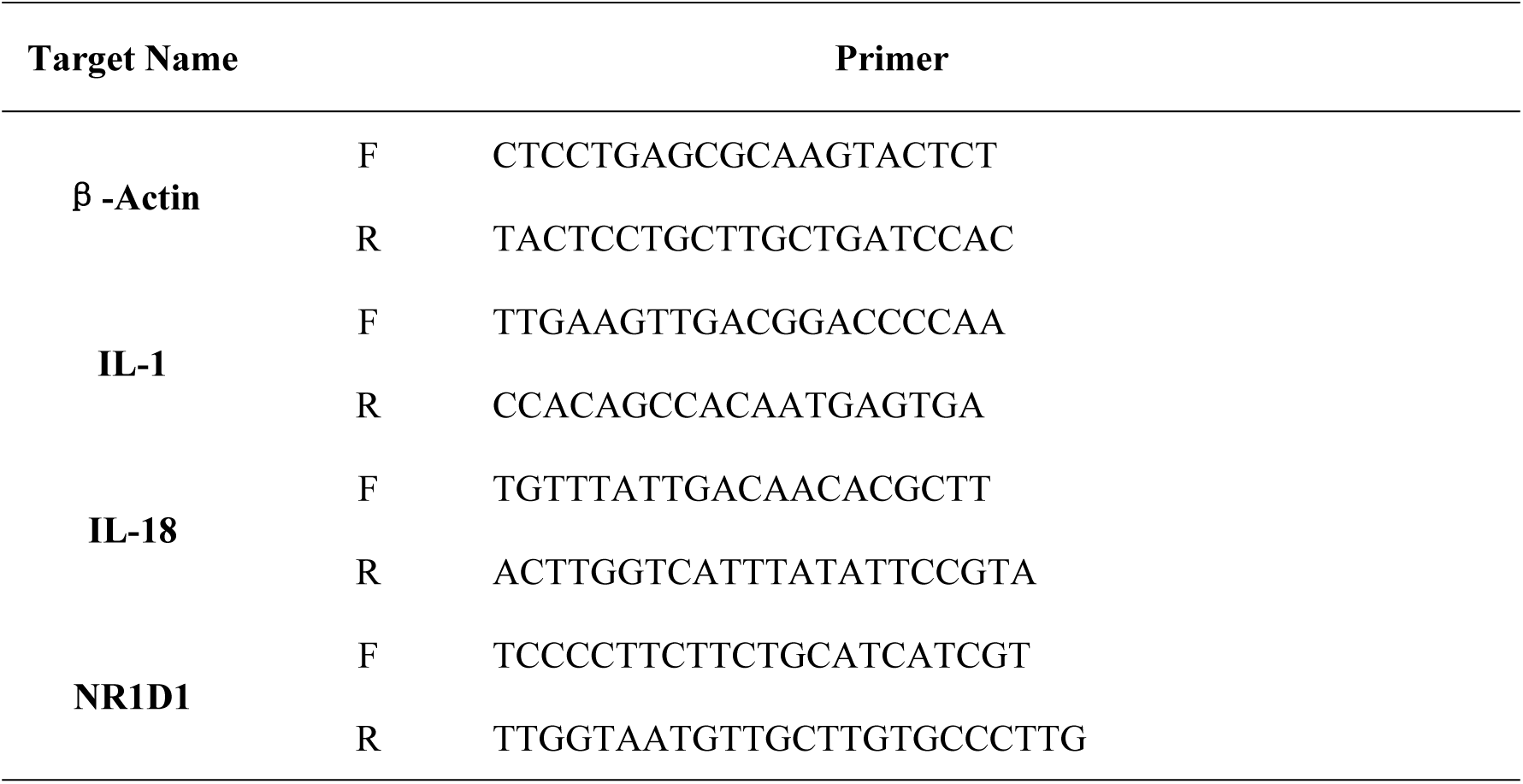
PCR Primer Design.

### 1.8 Statistical Analysis

Experimental data were analyzed using SPSS 25.0 software, with measurement data presented as mean ± standard deviation. Student’s t-test was employed for significance analysis between two groups, and one-way ANOVA was used for comparisons among multiple groups, with Dunnett’s t-test for inter-group differences. All statistical tests were two-sided, with the alpha set at 0.05. A p-value < 0.05 indicates statistical significance.

## Results

### 2.1 Establishment of Mouse Models and Changes in Physiological Indices and Blood Lipid Levels

After 12 weeks of high-fat feeding, the body weights of mice in the high-fat diet (GZ) group, UC group, and SR9009 group (JD) were significantly higher than those in the control group (DZ) (*P=*0.015, 0.000, 0.005), confirming the successful establishment of the obesity model [Table 3, Figure 1].

**Figure 1.**
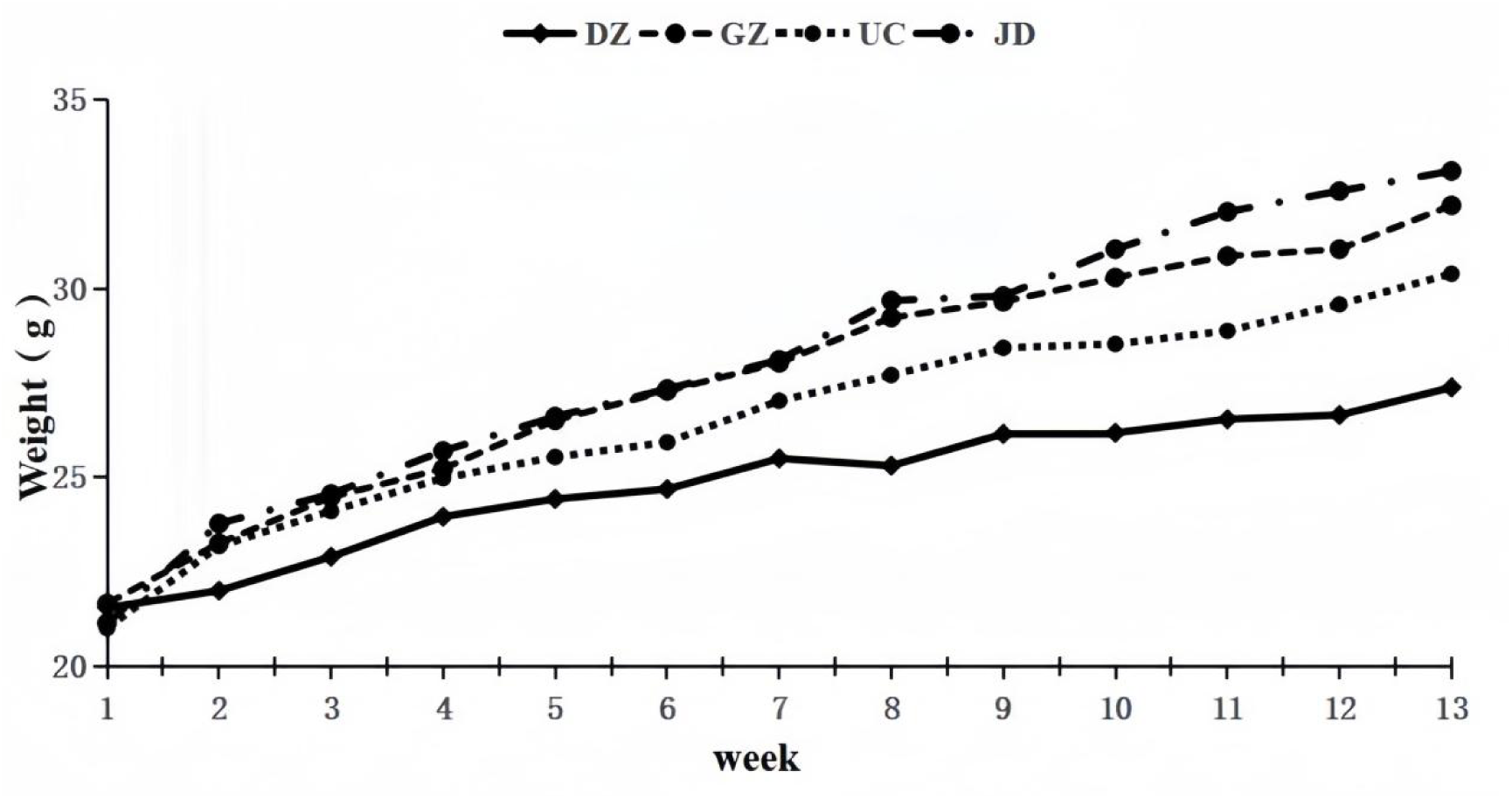
Changes in Body Weights of Mice Across Groups.

**Table 3.**
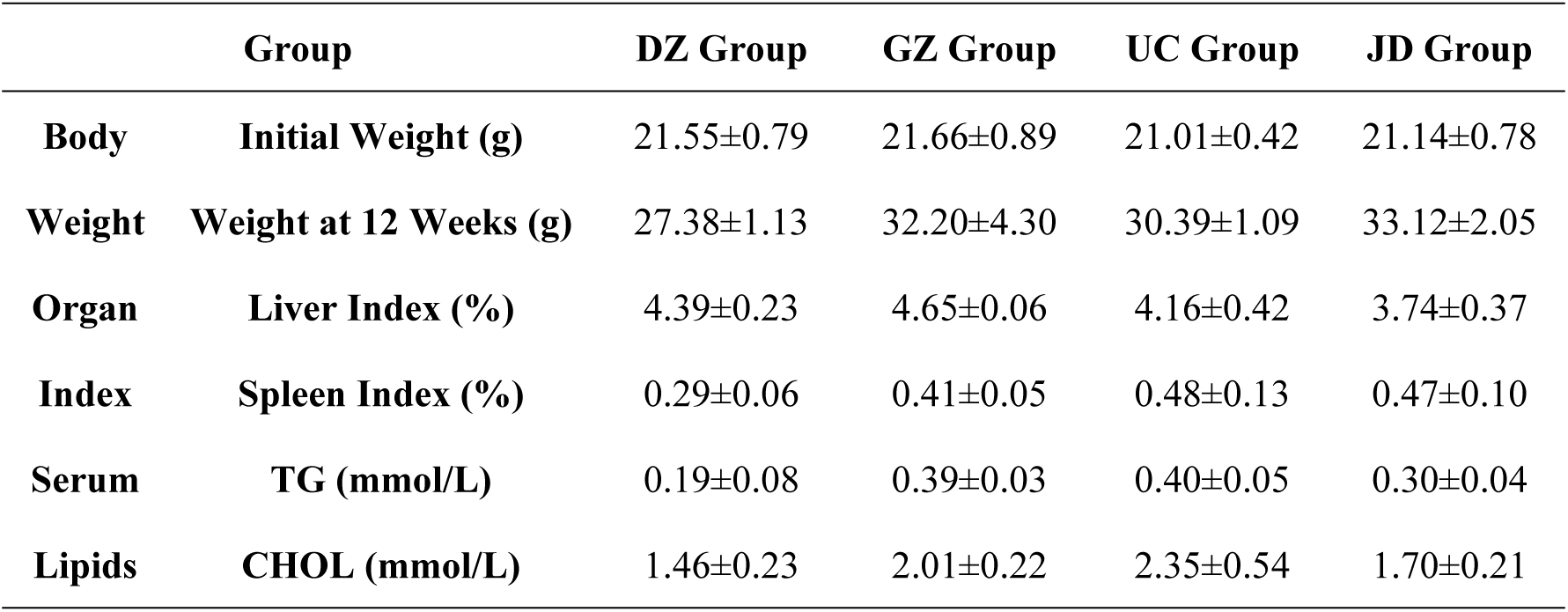
Body Weight, Organ Index, and Blood Lipid Levels in Mice.

In liver index results, the GZ group exhibited a higher liver index than the DZ group, though not reaching statistical significance (*P>*0.05). The UC and JD groups showed no significant differences in liver index compared to the DZ group (*P>*0.05) [Figure 2]. The spleen index indicated that the GZ and UC groups had significantly elevated spleen indices compared to the DZ group (*P=*0.001, 0.003) [Figure 2]. Compared to the UC group, the JD group’s liver index decreased by 10.10% following SR9009 intervention, with a slight decline in the spleen index, though both changes did not achieve statistical significance (*P>*0.05). These results indicate that DSS can induce visceral damage in mice, particularly evident by the significant increase in spleen index, consistent with manifestations in colitis models. Following SR9009 intervention, both liver and spleen indices showed varying degrees of reduction, suggesting a potential role for SR9009 in alleviating the inflammatory response of UC in mice.

**Figure 2.**
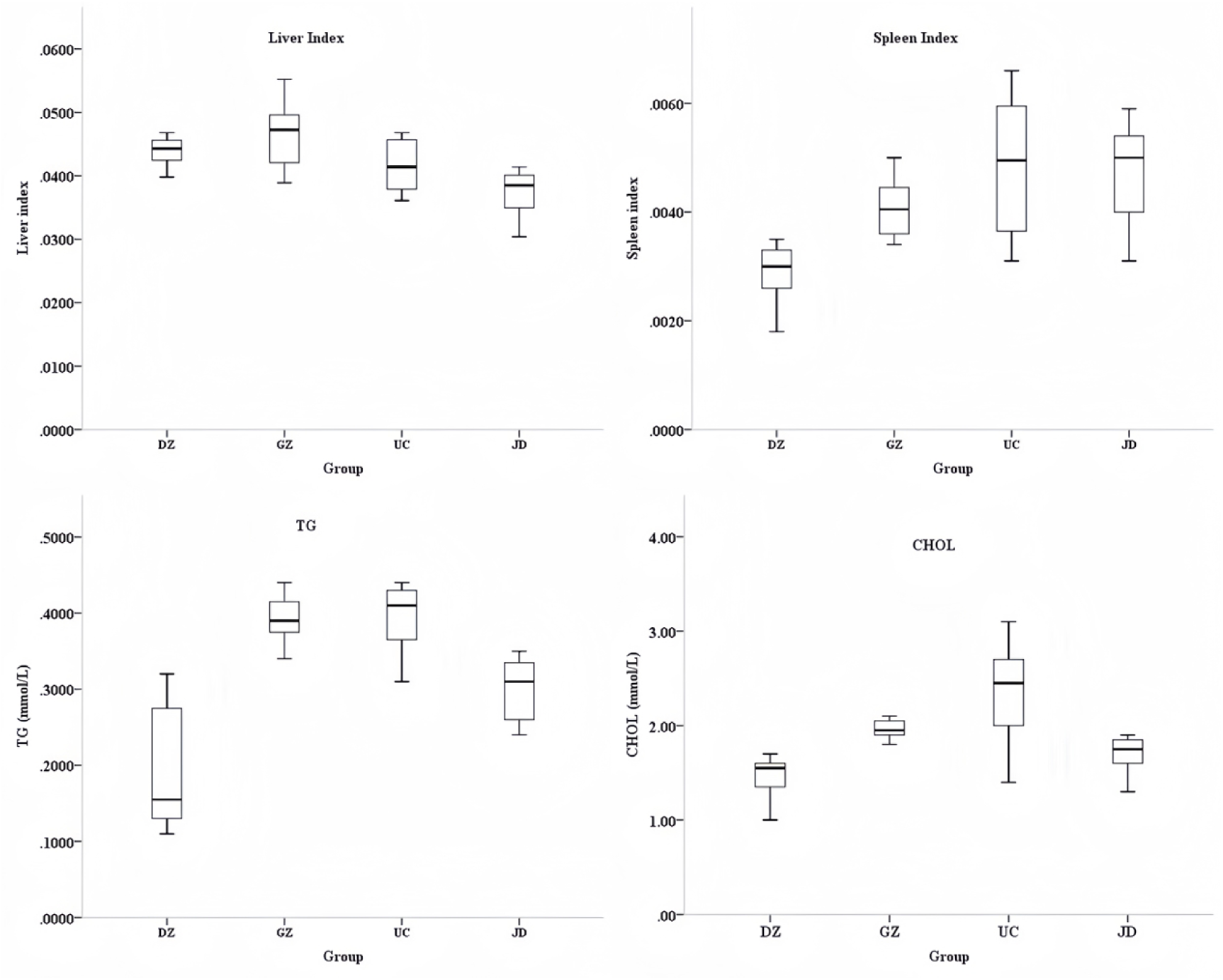
Visceral Indices and Blood Lipid Levels of Mice.

Compared with the DZ group, the serum TG and CHOL levels in the GZ group were significantly elevated (*P=*0.000, 0.000), confirming the successful establishment of the obesity model. The UC group also showed significantly increased serum TG and CHOL levels compared to the DZ group (*P=*0.000, 0.001). After SR9009 intervention, the JD group exhibited significantly reduced serum TG and CHOL levels compared to the UC group (*P=*0.001, 0.011), indicating SR9009’s role in ameliorating blood lipid abnormalities in obesity-associated UC mice [Figure 2].

### 2.2 Pathological Changes in Colonic Tissues

Histological examination of colonic tissues via HE staining assessed the extent of pathological damage (Figure 3). Mice in the DZ group exhibited intact colonic walls with evenly distributed crypt structures containing abundant goblet cells and no notable inflammatory cell infiltration. In contrast, the GZ group displayed relatively intact colonic walls with some inflammatory cell infiltration, yet evident crypt and glandular structures remained. In the UC group, significant degeneration and necrosis of colonic epithelial cells were observed, with destruction of crypt structures and a marked reduction in goblet cells, leading to extensive infiltration of inflammatory cells within the mucosa. Following SR9009 intervention, the JD group revealed partial destruction of crypt structures with minimal inflammatory cell infiltration and an increase in goblet cells. HAI scoring indicated that compared to the DZ group, both GZ and UC groups had significantly higher HAI scores (t=10.142, 14.836, *P=*0.000, 0.000). Conversely, the JD group showed significantly lower HAI scores compared to the UC group (t=4.854, *P=*0.000). These results indicate that DSS-induced colitis successfully established the UC mouse model, while SR9009 mitigated the colonic tissue damage attributable to UC.

**Figure 3.**
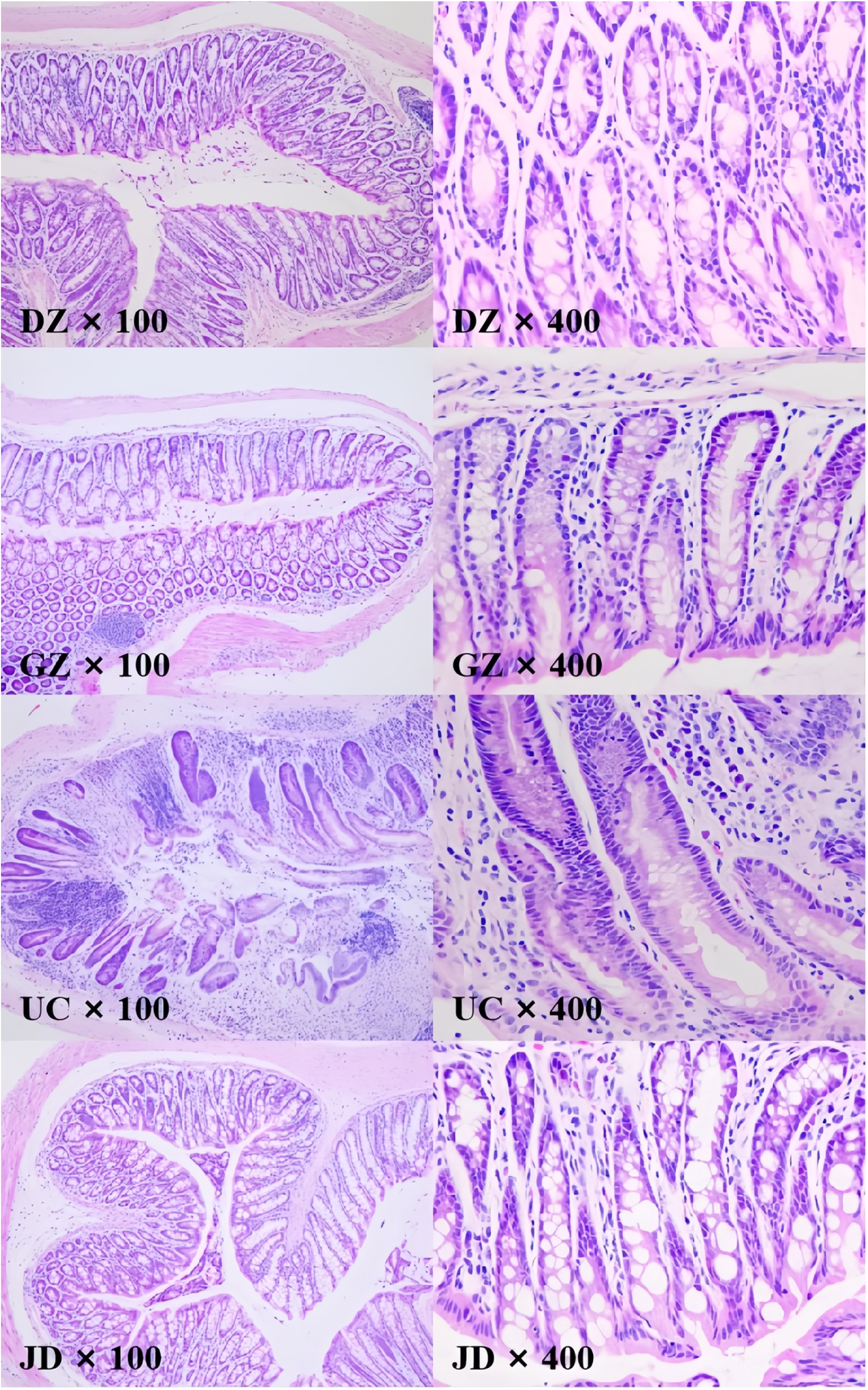
HE Staining of Mouse Colonic Tissues.

### 2.3 Changes in Serum Inflammatory Factor Levels

Serum samples were analyzed for IL-1β and IL-18 via ELISA (Figure 4). Compared to the DZ group, the GZ and UC groups demonstrated significantly elevated serum levels of the pro-inflammatory factor IL-1β (t=7.327, 20.201; *P=*0.000, 0.000), with similar trends observed for IL-18 levels (t=11.741, 24.480; *P=*0.000, 0.000). Following SR9009 intervention, the JD group exhibited significantly reduced serum IL-1β levels compared to the UC group (t=8.931, *P=*0.000), alongside a notable decrease in serum IL-18 levels (t=6.054, *P=*0.000). Data revealed that DSS induction significantly increased serum IL-1β levels from 271.27±18.73 pg/ml to 386.89±16.43 pg/ml and IL-18 levels from 356.58±15.64 pg/ml to 446.58±16.47 pg/ml in UC mice. Post-SR9009 treatment, serum IL-1β levels were reduced to 322.32±14.16 pg/ml and IL-18 levels to 386.50±24.80 pg/ml, indicating SR9009’s potential in alleviating inflammatory conditions in UC mice.

**Figure 4.**
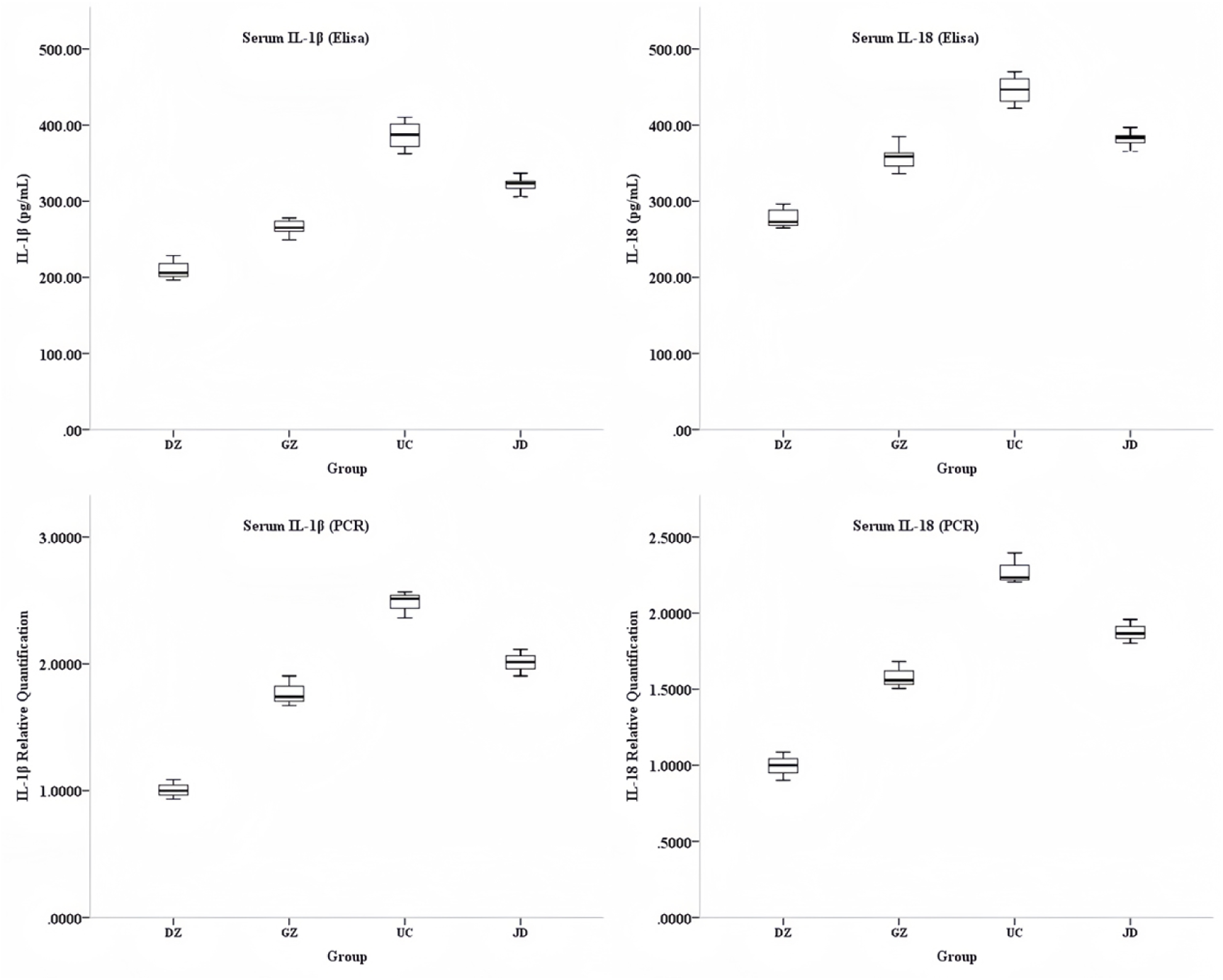
Expression Levels of IL-1β and IL-18 in Mouse Serum and Colonic Tissues.

We further assessed IL-1β and IL-18 expression levels in serum and colonic tissues using PCR, revealing that levels in the GZ and UC groups were significantly elevated compared to the DZ group (t=9.266, 19.442; *P=*0.001, 0.000), with similar increases in serum IL-18 (t=7.824, 16.046; *P=*0.001, 0.000). Post-SR9009 intervention, the JD group presented significantly lowered serum IL-1β levels compared to the UC group (t=5.461, *P=*0.005), with a similar significant decline in serum IL-18 levels (t=5.385, *P=*0.006). Additionally, IL-1β and IL-18 expression levels in colonic tissues were markedly elevated in the GZ and UC groups compared to the DZ group (*P=*0.000, 0.000; *P=*0.000, 0.000). Following SR9009 treatment, the JD group exhibited significant reductions in IL-1β and IL-18 expression levels in colonic tissues compared to the UC group (*P=*0.004, 0.003, Figure 4). These findings further validate the successful establishment of the DSS-induced UC mouse model and suggest that SR9009 exerts a therapeutic effect on the inflammatory response in UC.

### 2.4 Changes in Macrophages in Colonic Tissues

Flow cytometry was utilized to detect changes in macrophages within colonic tissues (Figure 5). Results indicated that the proportions of CD68-positive cells in the GZ and UC groups rose significantly from 5.84±0.35% to 16.32±0.34% and 34.44±0.27%, respectively, when compared to the DZ group (t=37.430, 111.247; *P=*0.000, 0.000). Following the SR9009 intervention, the JD group exhibited a reduced proportion of CD68-positive cells (23.01±0.19%) compared to the UC group, with significant differences observed (t=59.782, *P=*0.000). Conversely, the proportions of CD206-positive cells in the GZ and UC groups diminished compared to the DZ group, with CD206 levels declining from 30.01±0.66% to 15.42±0.17% and 4.98±0.34%, respectively (t=36.977, 58.499; *P=*0.000, 0.000). Post-SR9009 treatment, the JD group showed a significant increase in CD206-positive cell proportions compared to the UC group (t=15.982, *P=*0.000). Comprehensive analysis of CD68 and CD206 expression revealed a decline in the CD206/CD68 ratio in the GZ and UC groups, whereas the JD group exhibited an increased ratio compared to the UC group, with significant differences noted (*P<* 0.001). M1 macrophages are characterized by the high expression of CD68 and other protein markers, while M2 macrophages are distinguished by elevated expression of CD206. Based on these findings, DSS induction in obese mice promotes a shift towards M1-type macrophage polarization while inhibiting the transition towards M2 macrophages. In contrast, SR9009 treatment appears to modulate macrophage polarization by inhibiting M1 polarization and promoting M2 polarization, thus enhancing intestinal mucosal immune regulation and alleviating gut inflammation.

**Figure 5.**
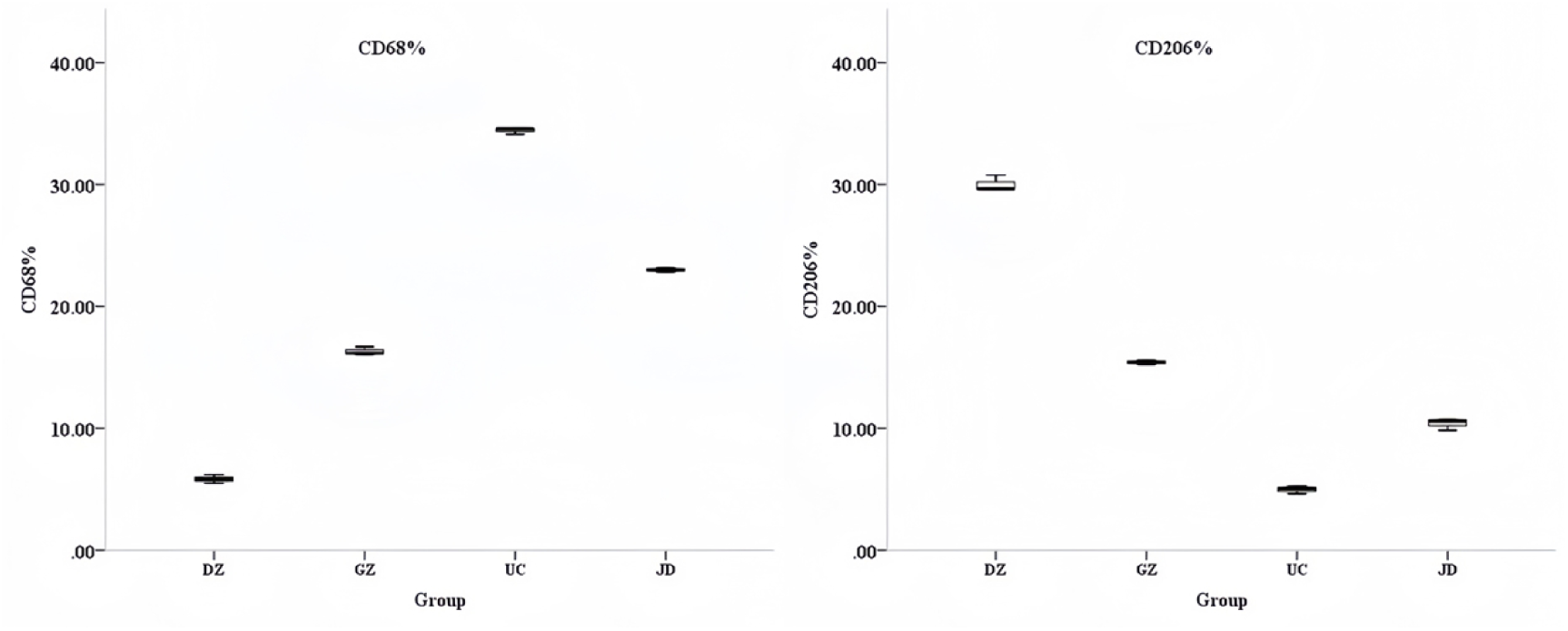
Changes in Macrophages in Mouse Colonic Tissues.

### 2.5 Expression Levels of Circadian Rhythm Factor NR1D1

To assess the expression of the circadian rhythm factor NR1D1, its levels in serum and colonic tissues were monitored (Figure 6). Western blot results indicated that NR1D1 protein expression in the colonic tissues of the GZ and UC groups was significantly reduced compared to the DZ group, whereas the JD group exhibited a notable increase in NR1D1 protein expression relative to the UC group. Further PCR analysis revealed that NR1D1 mRNA levels were significantly downregulated in the serum of the GZ and UC groups compared to the DZ group, with the UC group exhibiting even lower levels (t=3.850, 8.612; *P=*0.018, 0.001). Post-SR9009 intervention, serum NR1D1 mRNA levels in the JD group were significantly elevated in comparison to the UC group (t=3.850, *P=*0.004). In colonic tissue analyses, NR1D1 mRNA levels were also significantly decreased in the GZ and UC groups compared to the DZ group, with the UC group presenting lower levels (t=4.550, 7.782; *P=*0.010, 0.001). Following SR9009 treatment, NR1D1 mRNA levels in colonic tissues were significantly upregulated in the JD group compared to the UC group (t=5.142, *P=*0.007). Collectively, these results indicate that NR1D1 expression is downregulated in the serum and colonic tissues of UC patients and that SR9009 intervention may alleviate colonic inflammation by upregulating NR1D1 levels.

**Figure 6.**
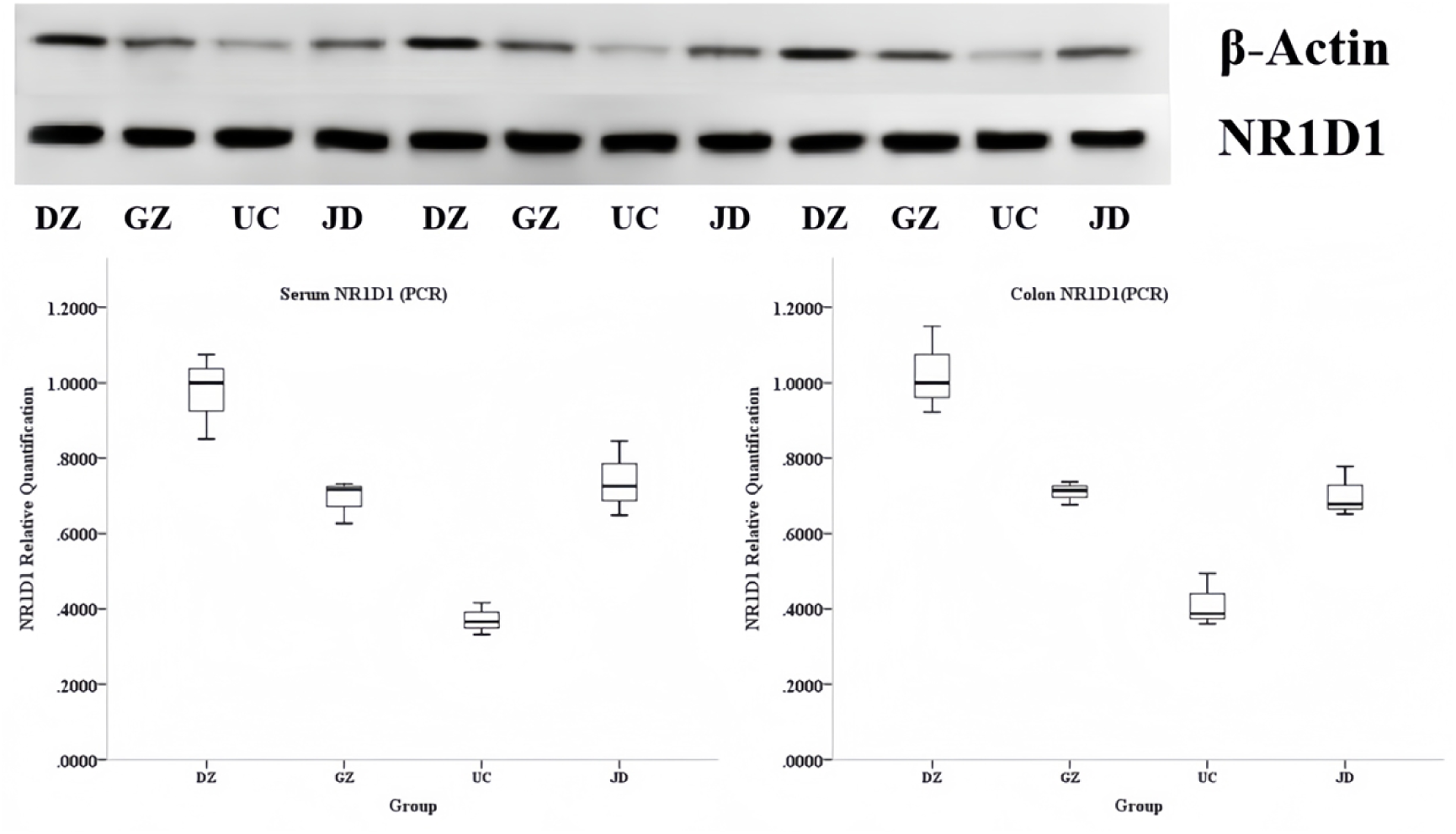
Expression of NR1D1 in Mouse Serum and Colonic Tissues.

## Discussion

Globally, the incidence rates of obesity and ulcerative colitis (UC) are significantly rising, progressively becoming a pressing public health issue. Obesity, as a complex chronic disease, is characterized not only by excessive fat accumulation in the body but also reflects a series of metabolic disorders. UC, on the other hand, is a chronic nonspecific inflammatory bowel disease with an incompletely understood etiology, generally believed to result from the interplay of various factors, including genetics, environment, immune responses, and gut microbiota. In recent years, with changes in lifestyle and dietary structures, obesity has become prevalent, and inflammatory bowel disease is increasingly diagnosed in obese individuals [33–35]. Some studies suggest that obesity is considered a risk factor in the pathogenesis of UC, with the incidence of UC increasing globally, particularly in emerging industrialized countries and regions [36,37]. A retrospective study conducted in 2021 found that 20.6% of patients with ulcerative colitis were obese, and 34.8% were overweight, although the impact of obesity on UC remains uncertain [38]. While obesity and ulcerative colitis may appear to be two distinct diseases, a growing body of research indicates a close association between the two. On one hand, obesity may influence the risk, progression, and treatment outcomes of ulcerative colitis through various mechanisms. Factors such as chronic inflammatory states, gut dysbiosis, and metabolic disturbances caused by obesity may create conditions conducive to the onset and progression of UC. Conversely, patients with ulcerative colitis may experience increased risks of obesity due to malabsorption issues arising from the disease, gut dysfunction, and certain medications used during treatment. A thorough exploration of the relationship between obesity and ulcerative colitis is crucial for elucidating the pathogenesis of obesity-related UC, developing precise prevention and treatment strategies, and improving patient outcomes [39–41].

In this study, we successfully established a mouse model of obesity through a high-fat diet and subsequently induced UC using DSS. The high-fat diet not only resulted in increased body weight in the mice but also triggered dysregulation of lipid metabolism, leading to elevated levels of serum TG and CHOL, further confirming the successful establishment of the obesity model [42]. Changes in liver and spleen indices reflect the health status and inflammatory response of the visceral organs in mice. This study observed only a trend of increasing liver indices following high-fat diet feeding, suggesting that prolonged high-fat diets may lead to fat accumulation in the liver and subsequent morphological and functional changes, although these changes did not reach statistical significance, possibly due to factors such as experimental duration, sample size, or others. Regarding spleen indices, both the high-fat diet group and the UC model group exhibited significantly higher spleen indices compared to the control group, indicating that both high-fat diets and DSS induction contributed to spleen enlargement, likely related to the activation of inflammatory responses. The spleen, as a vital immune organ, activates immune cells in response to inflammation, resulting in increased spleen volume. DSS-induced UC may trigger systemic or localized inflammatory responses, causing the spleen to participate in immune defense, thereby increasing spleen indices [43–45]. Histological examination of colonic tissues stained with HE and HAI scoring revealed partial inflammatory cell infiltration in the colonic walls of high-fat diet mice, whereas mice with obesity-associated ulcerative colitis exhibited severe damage to the mucosal epithelial layer, destruction of crypt structures, and significant reductions in goblet cells, severely compromising intestinal function and leading to symptoms such as diarrhea, abdominal pain, and hematochezia [39,46].

IL-1β and IL-18 are critical pro-inflammatory factors playing key roles in inflammatory responses; they can activate immune cells to promote the release of inflammatory mediators, leading to inflammation’s onset and progression. This study found that high-fat diets might alter the gut microbiota environment, affecting the functionality of intestinal immune cells and promoting the production of IL-1β and IL-18. DSS-induced UC further exacerbated this inflammatory response, substantially elevating serum levels of IL-1β and IL-18. PCR results similarly indicated increased synthesis of IL-1β and IL-18 at the gene level, likely associated with upregulation of relevant gene expressions [47]. Macrophages, as essential components of the immune system, play a crucial role in maintaining immune balance and responding to pathogen invasions. In the context of intestinal inflammation, the polarization state of macrophages changes, closely associated with the onset, progression, and resolution of inflammation. CD68 is a hallmark protein of M1 macrophages, and this study revealed that under a high-fat diet and DSS induction, the increase in CD68-positive rates indicates a trend toward enhanced M1 macrophage polarization in colonic tissues. This shift may arise from disturbances in gut microbial balance caused by high-fat diets and DSS, leading to increased pathogen or damage-associated molecular pattern signals that stimulate macrophages towards M1 polarization. M1 macrophages possess potent pro-inflammatory capabilities, secreting significant amounts of pro-inflammatory factors such as IL-1β, IL-6, and TNF-α, which further exacerbate intestinal inflammation, causing tissue damage and dysfunction [48]. Conversely, the expression of CD206, a marker for M2 macrophages, is reduced, indicating suppressed M2 polarization. M2 macrophages have anti-inflammatory and tissue repair functions, releasing anti-inflammatory factors such as IL-10 and TGF-β that promote the resolution of inflammation and tissue repair. In this study, the suppression of M2 macrophage polarization diminished the ability to counteract inflammation, further aggravating intestinal inflammatory responses [49]. The overall analysis of both expressions indicates a relative increase in M1 macrophages and a decrease in M2 macrophages in the colonic tissues, shifting the balance of macrophage polarization towards pro-inflammatory directions and intensifying intestinal inflammation.

NR1D1, also known as Rev-erbα, is encoded by a gene located on chromosome 17q21.31, containing multiple functional domains. Its N-terminal comprises a highly variable transcriptional activation domain (AF-1) that interacts with other transcription factors to regulate gene transcription activity [50]. NR1D1 was initially identified as a core component of the circadian clock regulatory network, playing a key role in maintaining circadian rhythms. It forms negative feedback regulatory loops with clock genes such as Bmal1, precisely orchestrating the rhythmic expression of circadian genes and influencing various physiological processes, including sleep-wake cycles, hormone secretion, and metabolism [51,52]. Additionally, NR1D1 participates in regulating critical metabolic processes, such as lipid and glucose metabolism [3]. NR1D1 can directly bind to the promoter regions of inflammatory cytokine genes, such as IL-6 and IL-1β, inhibiting their transcription to exert anti-inflammatory effects [53–55]. This study observed a significant downregulation of NR1D1 protein expression in the colonic tissues of the high-fat diet and UC groups, indicating that NR1D1 expression is suppressed in the pathological states of obesity and UC. In obesity, metabolic dysregulation may affect the regulatory mechanisms of circadian genes, leading to decreased NR1D1 expression. DSS-induced UC exacerbates this downregulation, likely due to inflammatory responses interfering with intracellular signaling pathways, further impacting NR1D1 gene transcription and translation processes. PCR results reaffirm that both obesity and UC lead to downregulation of NR1D1 expression, with the effects being more pronounced in UC. Multiple factors may synergistically contribute to the transcriptional suppression of NR1D1, such as methylation modifications in the promoter region and aberrant transcription factor binding in the pathogenesis of obesity and UC.

The circadian rhythm factor NR1D1 is indispensable for maintaining normal physiological rhythms and internal stability, and its role in inflammatory responses has garnered increasing attention. In 2012, Solt LA et al. synthesized two NR1D1 ligands, SR9009 and SR9011, confirming their capacity to enhance NR1D1 transcriptional function [56]. SR9009, as a highly selective REV-ERBα/β agonist, may activate downstream signaling pathways by binding to REV-ERBα/β, promoting NR1D1 gene transcription and translation, thereby increasing NR1D1 protein expression, which helps restore normal circadian rhythms, regulate related gene expressions, and reduce inflammation. Supporting our results, we found that following SR9009 intervention in the ulcerative colitis mouse model, both liver and spleen indices decreased to varying degrees. Although these changes did not achieve statistical significance, they nonetheless suggest that SR9009 may have protective effects against visceral inflammatory responses in mice. SR9009 may modulate metabolic and immune functions, reducing the release of inflammatory factors, and thus providing some protection to the liver and spleen. Following SR9009 intervention, pathological conditions in the colonic tissues of ulcerative colitis mice showed marked improvement, with reduced inflammatory cell infiltration and increased goblet cells, indicating that SR9009 could inhibit the recruitment of inflammatory cells to colonic tissues while promoting goblet cell proliferation or differentiation, thereby aiding in the repair of damaged intestinal mucosal barriers and maintaining the normal structure and function of colonic tissues. This study suggests that SR9009 may exert anti-inflammatory effects through multiple mechanisms, including inhibiting the activation of inflammatory signaling pathways, thereby reducing transcription and translation processes of pro-inflammatory factors such as IL-1β and IL-18. SR9009 effectively lowered serum inflammatory factor levels and significantly reduced IL-1β and IL-18 expression in colonic tissues, thus alleviating inflammatory damage. High expression of IL-1β and IL-18 in colonic tissues leads to inflammatory cell infiltration and tissue damage, while SR9009’s inhibition of their expression helps maintain the normal structure and function of colonic tissues. Furthermore, SR9009 appears to regulate intestinal immune cell functions, reducing inflammatory cell accumulation in colonic tissues, and thereby decreasing IL-1β and IL-18 levels. Moreover, SR9009 may enhance the repair and regeneration of colonic tissues, bolstering the functionality of the intestinal mucosal barrier and mitigating the impact of inflammatory factors on colonic tissues [57–60]. Additionally, SR9009 may inhibit macrophage polarization towards M1 and promote polarization towards M2, thereby decreasing M1 macrophage numbers while increasing M2 macrophage counts, reducing pro-inflammatory factor secretion, ameliorating inflammatory responses, and enhancing anti-inflammatory and tissue repair capacities. SR9009 could inhibit NF-κB pathway activation, reducing the transcription of pro-inflammatory genes, thus lowering the expression of M1 macrophage markers such as CD68, while simultaneously activating the PPAR-γ signaling pathway and upregulating M2 macrophage markers such as CD206 [61,62]. Collectively, these findings indicate that SR9009 intervention enhances intestinal inflammatory responses through the upregulation of NR1D1 levels.

As an NR1D1 agonist, SR9009 plays a critical role in inflammation suppression, potentially through regulatory mechanisms at multiple levels. At the gene transcription level, SR9009 may interact with NR1D1, where NF-κB serves as a key transcription factor regulating the transcription of IL-1β and IL-18 genes. Inhibition of NF-κB signaling pathways can suppress their activation, consequently reducing the transcription of IL-1β and IL-18 genes and their synthesis. At the protein translation level, SR9009 may influence mRNA stability or the activity of translation initiation factors, affecting the translation processes of IL-1β and IL-18. SR9009 may interact with mRNA-binding proteins or translation initiation factors, reducing the stability of IL-1β and IL-18 mRNA, facilitating its degradation, or inhibiting the activity of translation initiation factors, obstructing ribosomal binding to mRNA, thus reducing IL-1β and IL-18 translation and their protein levels. Furthermore, SR9009 may also regulate the activity of immune cells such as T cells, B cells, and macrophages, inhibiting their secretion of IL-1β and IL-18 [24,63]. Modulating the polarization state of macrophages is also a vital mechanism through which SR9009 suppresses colonic inflammation. During macrophage polarization, REV-ERBα/β may regulate the activity of signaling pathways such as NF-κB and JAK-STAT, influencing the direction of macrophage polarization. The NF-κB signaling pathway plays a crucial role in M1 macrophage polarization. When macrophages encounter pathogens or inflammatory stimuli, the NF-κB signaling pathway is activated, leading to upregulation of M1 macrophage-associated gene expression and promoting macrophage polarization towards M1. Conversely, the activation of REV-ERBα/β can inhibit NF-κB pathway activation. Studies have indicated that REV-ERBα/β can interact with the NF-κB subunit p65, inhibiting its nuclear translocation and thereby blocking NF-κB pathway activation. The JAK-STAT signaling pathway also participates in the regulation of macrophage polarization, with Th1 cytokines such as IFN-γ activating the JAK-STAT1 pathway to promote M1 polarization, whereas Th2 cytokines like IL-4 and IL-13 activate the JAK-STAT6 pathway to encourage M2 polarization, thereby exerting anti-inflammatory and tissue repair effects [64–67]. The intervention of SR9009 can activate Rev-erbα, enabling it to bind to RORE elements in the NR1D1 gene promoter region, recruiting co-repressor factors, and inhibiting NR1D1 gene transcription, thus upregulating NR1D1 expression. This process restores normal NR1D1 expression in the body and maintains circadian rhythm stability. In addition to directly regulating NR1D1 expression, SR9009 may also influence the expression of other circadian rhythm-related genes and pathways. Within the core feedback loop of the molecular clock, BMAL1 and CLOCK form heterodimers that bind to E-box elements in downstream gene promoters, promoting the transcription of a series of circadian rhythm-related genes, including NR1D1. NR1D1 and other nuclear receptors like RORα bind to RORE elements, thereby inhibiting and activating BMAL1 expression, respectively, forming a complex regulatory network [68]. In the pathological processes of obesity and ulcerative colitis, this regulatory network may be disrupted, leading to circadian rhythm disorders. After SR9009 activates Rev-erbα, it may adjust the expression of other genes within this regulatory network to restore normal circadian rhythm regulation. For instance, SR9009 may inhibit BMAL1 expression, reducing its activation effect on downstream genes, thus adjusting the expression patterns of circadian rhythm-related genes. Moreover, SR9009 may also affect the functions of other nuclear receptors such as RORα, indirectly influencing immune cell functions and inflammatory responses, thereby alleviating symptoms of obesity-associated ulcerative colitis [69]. While this study has explored the mechanisms of SR9009, there are still some limitations that require further investigation in future research. In terms of the mechanisms of anti-inflammatory response, while we have revealed that SR9009 exerts anti-inflammatory effects by inhibiting the NF-κB signaling pathway and modulating macrophage polarization, inflammation is a highly complex biological process involving interactions among various cell types and signaling pathways. It remains to be seen whether SR9009 also regulates the inflammatory response through other unknown mechanisms (such as gut microbiota, neuroendocrine systems, etc.), and how these mechanisms interact with one another. In terms of circadian rhythm regulation, future studies should observe the effects of SR9009 on the expression of circadian rhythm-related genes and proteins, as well as its impact on the physiological functions of mice and improvements in disease symptoms, by altering environmental factors such as light cycles and temperature or by reducing body weight [70]. Future research should focus on optimizing experimental models, expanding detection indicators, and delving deeper into the mechanisms to comprehensively reveal the pathogenesis of obesity-associated ulcerative colitis and the mechanisms of SR9009, thereby establishing a solid theoretical basis for developing more effective treatment strategies.

## Conclusion

This study systematically explored the effects and potential mechanisms of SR9009 on a mouse model of obesity-associated ulcerative colitis. SR9009 alleviates symptoms related to obesity-associated UC by upregulating NR1D1 expression and modulating circadian rhythm-related genes and pathways, thereby restoring normal circadian regulation. This finding provides new insights into the understanding of SR9009’s mechanisms of action and offers potential therapeutic targets and strategies for treating obesity-associated ulcerative colitis and other conditions related to circadian rhythm disruptions. Future research may further investigate the specific molecular mechanisms by which SR9009 regulates circadian rhythms as well as the intrinsic connections between circadian rhythm disorders and diseases such as obesity and ulcerative colitis, laying the groundwork for developing more effective treatments and advancing the field of therapy for obesity-associated ulcerative colitis.

## Declarations

### Ethics approval and consent to participate

This study was approved by the Ethics Committee of the Beijing Jishuitan Hospital, Capital Medical University. All the methods were carried out according to relevant guidelines and regulations.

### Authors’ contributions

Zhixue Zheng conceived and designed the experiments, analyzed and interpreted the data, and wrote the manuscript, Xuan Cai and Yaqi Liu analyzed the data, Jingtao Bi revised the manuscript. All authors reviewed the manuscript. All authors agreed to publish this article.

### Consent for publication

All authors have consented to the publication of this manuscript.

### Availability of data and material

All data generated or analyzed during this study are included in the article and its supplementary materials.

